# Opposing forms of adaptation in mouse visual cortex are controlled by distinct inhibitory microcircuits and gated by locomotion

**DOI:** 10.1101/2020.01.16.909788

**Authors:** Tristan G. Heintz, Antonio J. Hinojosa, Leon Lagnado

**Affiliations:** Sussex Neuroscience, School of Life Sciences, University of Sussex, Brighton BN1 9QG, UK

## Abstract

Cortical processing of sensory signals adjusts to changes in both the external world and the internal state of the animal. We investigated the neural circuitry by which these processes interact in the primary visual cortex of mice. An increase in contrast caused as many pyramidal cells (PCs) to sensitize as depress, reflecting the dynamics of adaptation in different types of interneuron (PV, SST and VIP). Optogenetic manipulations demonstrate that the net effect within PCs reflects the balance of PV inputs, driving depression, and a subset of SST interneurons, driving sensitization. Locomotor behaviour increased the gain of PC responses by disinhibition through both the VIP->SST and SST->PV pathways, thereby maintaining the balance between opposing forms of plasticity. These experiments reveal how inhibitory microcircuits interact to purpose different subsets of PCs for the signalling of increases or decreases in contrast while also allowing for behavioural control of gain across the population.

## Introduction

The sensory world is dynamic and so are the properties of the neural circuits that process the information it provides (Weber et al., 2019). Adjustments in these circuits driven by the recent history of stimulation are generally termed adaptation, and are thought to improve the efficiency with which information can be extracted in a changing sensory environment (Whitmire and Stanley, 2016). In the mammalian cortex, the processing of external stimuli is also modulated by changes in the internal state of the animal, reflected in behaviours such as the sleep-wake cycle, locomotion and arousal (Harris and Thiele, 2011; Ferguson and Cardin, 2020). How do endogenous changes in cortical state interact with adaptive responses to the external world? To investigate this question, we have characterised the circuitry underlying contrast adaptation in primary visual cortex (V1) and the impact of locomotor behaviour on that circuitry (Fu et al., 2014).

The most commonly observed form of adaptation in the visual system is a decrease in sensitivity to a constant feature of the input. This feature might be the mean luminance or contrast, or statistical properties of higher-order, such as spatial patterns or movements in particular directions (Baccus and Meister, 2002; Chung et al., 2002; Kohn and Movshon, 2003; Levy et al., 2013; Solomon and Kohn, 2014; Whitmire and Stanley, 2016). Adaptation to contrast as a depressing response has been observed throughout the visual system, beginning in the synaptic output of retinal bipolar cells (Ozuysal and Baccus, 2012; Nikolaev et al., 2013), through retinal ganglion cells (Smirnakis et al., 1997), the dorsal lateral geniculate nucleus (dLGN) (Solomon et al., 2004; King et al., 2016) and V1 (Benucci et al., 2013; Bonin et al., 2006; Dhruv and Carandini, 2014; King et al., 2016; Wilson et al., 2012). But adaptation in V1 is not simply inherited from the retina: local processing generates time-dependent changes in gain of PCs, although the circuit mechanisms remain unclear (Dhruv and Carandini, 2014; King et al., 2016; Keller et al., 2017).

Inhibitory interneurons play a key role in controlling the gain and dynamics of signals in sensory cortex (reviewed in Ferguson and Cardin, 2020). There are many types of interneuron but the use of Cre-driver mouse lines has provided a way of separating the large majority of these into three broad groups, expressing one of Parvalbumin (PV), Somatostatin (SST) or Vasoactive Intestinal Polypeptide (VIP) (Tremblay et al., 2016). The connections between these and pyramidal cells (PCs) have been studied intensively (Pfeffer et al., 2013; Pi et al., 2013; Yetman et al., 2019) and have revealed some of the mechanism by which sensitivity in V1 is modulated in response to changes in internal state, in particular the dramatic increase in sensitivity synchronized to locomotor behaviour. Cholinergic inputs from the basal forebrain activated during running excite VIP interneurons which results in excitation of PCs through a VIP->SST disinhibition pathway (Fu et al., 2014; Niell and Stryker, 2010; Pakan et al., 2016; Dipoppa et al., 2018). We have less understanding of the role of inhibitory circuits in controlling adaptation to a visual stimulus (Harris and Shepherd, 2015; King et al., 2016), in part because most physiological studies of sensory cortex have used anaesthetized animals in which inhibitory mechanisms are defective, altering fundamental aspects of circuit function including adaptation (Haider et al., 2013; Keller et al., 2017).

We find that PCs in V1 of awake mice experience opposing forms of plasticity: while some respond to an increase in contrast with high initial gain and then depress over periods of seconds, as many respond with low initial gain but then gradually sensitize so that the average activity across the population is relatively constant over time. Adaptation over a similar time-course occurs in interneurons but while VIP- and PV-positive cells sensitize, SST-interneurons also show opposing forms of plasticity. To understand the circuits underlying these varying dynamics we used optogenetics to activate or inhibit the major classes of interneuron and this revealed two levels of time-dependent gain-control in PCs. The first level reflects the balance of *direct* inputs received from PV interneurons driving depression and SST interneurons driving sensitization. These inputs are in turn modulated through two disinhibition pathways, VIP->SST and SST->PV, both of which become engaged during locomotion, thereby increasing the gain across the population of PCs while also maintaining the balance between depressing and sensitizing forms of adaptation. This study outlines some of the basic mechanisms by which adaptation to external stimuli and changes in behavioural state interact to control gain within the visual cortex.

## Results

### Opposing forms of adaptation across the population of pyramidal cells

To investigate the local circuit mechanisms contributing to contrast adaptation in V1, we used awake mice in which inhibition was intact (Haider et al., 2013). Neurons expressing the calcium reporter GCaMP6f were imaged in Layer 2/3 and stimuli consisted of drifting sinusoidal gratings presented for 10 s (20° visual field, 80% contrast, 1 Hz, see Methods). The duration of the stimulus was chosen based on the similar time-scale of adaptive effects observed in the retina (Ozuysal and Baccus, 2012; Johnston et al., 2019) and, more recently, in V1 of awake mice (Keller et al., 2017). The responsivity of neurons in V1 is strongly dependent on locomotion (Niell and Stryker, 2010; Fu et al., 2014) so we began by confining our analysis to measurements made while the mouse was running on a trackball.

Exposure to the high-contrast stimulus caused a range of adaptive effects across the population of PCs, as shown in Fig. 1. Within a field of view (Fig. 1A), some neurons generated strong initial responses that then depressed gradually over the next 10 s, as shown in Fig. 1B (ROI numbers 0-50) and Fig. 1C (top). Other neurons generated weak initial responses but the gain then increased gradually over a similar time-course of seconds (Fig. 1B, ROI numbers >300 and Fig. 1C, bottom). These opposing forms of plasticity represented either end of a continuum in the center of which many PCs did not show any significant adaptation to contrast (Fig. 1B, ROI numbers 100-300, Fig. 1C, middle).

**Figure 1.**
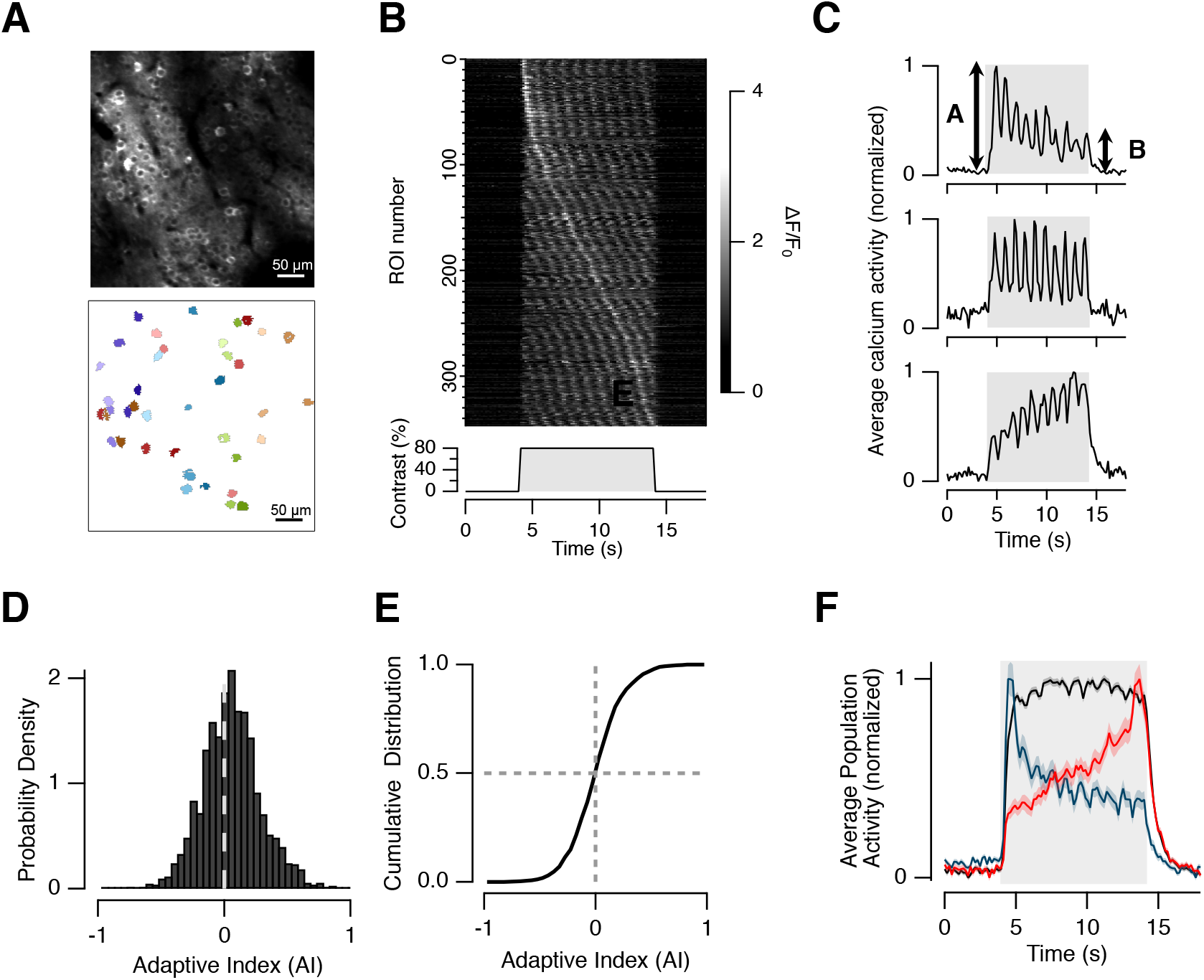
Opposing forms of plasticity in pyramidal cells. (**A**) An example of a field of view containing pyramidal cells (PCs) expressing GCaMP6f (top) and corresponding regions-of-interest (ROIs) defining individual neurons (see Methods). (**B**) Raster plot showing averaged responses from 350 PCs chosen at random from 10 experimental sessions in 3 mice. Neurons have been sorted according to time of peak response to the high-contrast stimulus: short delays correspond to depression (top) and longer delays to sensitization (bottom). (**C**) Average response from neurons 0-50 (top, depressing), 100-200 (middle, no significant adaptation) and 300-350 (bottom, sensitizing). For each neuron an Adaptive Index (AI) was calculated as (A-B)/(A+B) where A is the average response to the first two cycles of the 1 Hz stimulus and B the response to the last two cycles (top). (**D**) Distribution of the AI of 2441 pyramidal cells in 18 mice. (**E**) Cumulative distribution of AIs from measurements in D. The distribution of AIs in awake mice is symmetrical and centred around zero. **(F)** Normalized average response across all PCs from (D) and (E). The black trace is the population average superimposed on the of the 10% of neurons with the highest AI (blue) and 10% with the lowest (red). Note the similar time-course of depressing and sensitizing adaptation and the relatively constant activity averaged across the population of PCs.

To survey these adaptive effects across the population of PCs we quantified the strength and polarity of the stimulus-induced change in gain as an Adaptive Index (AI), defined as AI = (A−B)/(A+B), where A is the average response over the first two cycles of the 1 Hz stimulus and B is the average response over the last two cycles (Fig. 1C). Thus neurons that depress have an AI > 0, and those that depress an AI < 0. Surprisingly, as many PCs sensitized as depressed in response to the high-contrast stimulus: the distribution of AI across a population of 2441 neurons (n = 18 mice) was centered around zero (Fig. 1D and E). The time-course of the response averaged across all these is shown by the black trace in Fig. 1F, where it is compared with the top and bottom 10% of the AI distribution (blue and red, respectively). Notably, the activity of depressing and sensitizing PCs varied in a reciprocal manner, causing total activity to remain relatively constant. This behavior is qualitatively similar to “population homeostasis” observed in cat V1 during adaptation to orientated stimuli (Benucci et al., 2013). Experiments in anaesthetized animals have found that depressing adaptation is dominant in V1 (Keller and Martin, 2015; Sanchez-Vives et al., 2000), but these results demonstrate that there are two opposing forms of plasticity in awake mice and that these are in approximate balance across the population of PCs.

Why might it be the functional advantage of these different forms of adaptation? A decrease in gain is useful in that it prevents saturation, allowing for continued signalling of future increases in contrast (Kohn, 2007; Nikolaev et al., 2013; Solomon et al.; Wark et al., 2007). But such depressing adaptation comes at the cost of a reduced sensitivity to a future *decrease* in stimulus strength, at least when considering a single neuron. In the retina, a more complex picture has emerged: while some bipolar cells and ganglion cells depress after the initial response, others respond weakly at first but then sensitize (Kastner and Baccus, 2011; Nikolaev et al., 2013; Appleby and Manookin, 2019). The particular contribution of sensitizing neurons is to maintain sensitivity and signal future *decreases* in contrast more effectively, improving the overall rate of information transfer through the optic nerve when the contrast fluctuates (Kastner and Baccus, 2011; Appleby and Manookin, 2019). It may therefore be that opposing forms of plasticity in V1 also provide a strategy by which the population of PCs “hedge bets” in the face of an unpredictable future.

### Distinct modes of adaptation in different classes of interneuron

What are the circuit mechanisms that allow opposing gain changes to occur simultaneously within V1? A key role of local inhibition is suggested by the observation that optogenetic activation of all GABAergic neurons sufficient to “silence” PCs during a stimulus modifies subsequent adaptation (King et al., 2016). We hypothesized, therefore, that PCs showing increases and decreases in gain might have different connectivities with local inhibitory circuits. We began investigating this idea by assessing whether adaptive changes also occurred within different types of interneuron and found that they did. Fig. 2A shows averaged responses to the same high-contrast stimulus for the interneurons expressing VIP, SST or PV, with the corresponding distribution of AIs shown in Fig. 2B. Sensitization was the strongly dominant form of plasticity in both VIP and PV interneurons (average AI = −0.51 ± 0.032 and −0.26 ± 0.03; n = 96 and 89 cells, respectively), but SST interneurons were more heterogeneous, with 62% displaying net depression (AI > 0) and only 38% showing sensitization (average AI = 0.04 ± 0.02, n=152 cells; Fig. 2B). A distinct picture emerges, therefore, when interneurons are compared with PCs: contrast adaptation occurs in each class but depression and sensitization are *not* balanced.

**Figure 2.**
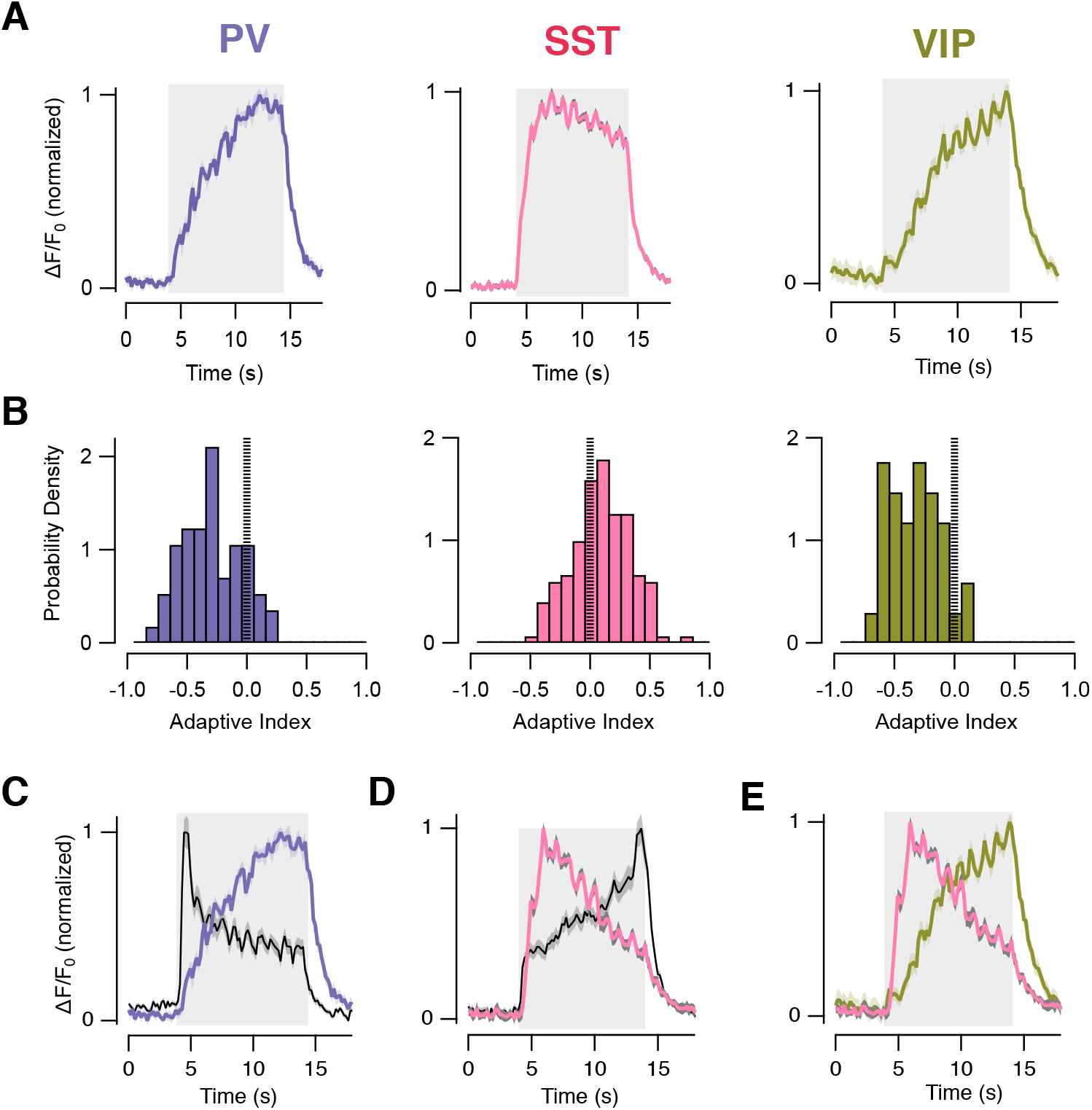
Distinct modes of adaptation in different classes of interneuron. **(A)** Average population response to prolonged high-contrast stimulus in PV (*left, purple trace*, n=1 mouse, 89 cells), SST (*Middle, pink trace; 3 mice,* 152 cells) and VIP interneurons (*right, green trace; 2 mice,* 96 cells) measured with GCaMP6f. The timing of the stimulus is shown by the grey bar. **(B)** Distribution of AI measured within each interneuron type shown in A. Note the strongly sensitizing responses in PV and VIP interneurons. SST cells were predominantly depressing but some were sensitizing. **(C)** Comparison of the dynamics of the response averaged in PV interneurons (*purple*) and the 10% of pyramidal cells with the highest AI (*black*). **(D)** Comparison of the dynamics of the response averaged in SST interneurons (*pink*) and the 10% of pyramidal cells with the lowest AI (*black*). **(E)** Comparison of the dynamics of the response averaged in SST (*pink*) and VIP (*green*) interneurons.

The time-course of adaptive effects in interneurons were similar to those observed in PCs, as would be expected if there was a causal relation. For instance, sensitization in PV cells roughly mirrored depression in PCs (Fig. 2C) while depression in SST interneurons with an AI > 0 mirrored sensitization in PCs (Fig. 2D). VIP interneurons make strong connections with SST interneurons (Pfeffer et al., 2013, Karnani et al., 2016), and, consistent with this, depression in SST interneurons with an AI > 0 followed a similar time-course to sensitization in VIP interneurons (Fig. 2E).

### PV interneurons drive depressing adaptation in pyramidal cells

Links between interneuron activity and adaptive effects in PCs were tested using optogenetics. The strongest inhibitory connections within Layer 2/3 are shown in Fig. 3A, based on electrical recordings, optogenetics and viral tracing (Pfeffer et al., 2013; Pi et al., 2013; Tremblay et al., 2016). We began by investigating the role of PV interneurons which are the most numerous inhibitory subtype and have been shown to modulate the gain of visual responses in PCs (Celio, 1986; Atallah et al., 2012). PV interneurons are also relatively isolated within inhibitory networks of the cortex, making strong and *direct* connections with PCs but not transmitting significantly to SST or VIP interneurons (Yetman et al., 2019), making it easier to interpret their role in adaptation.

First, we expressed both ChrimsonR and GCaMP6f in PV cells to assess how increased excitation altered the dynamics of their response to the high-contrast stimulus. ChrimsonR is a red-shifted form of channelrhodopsin that can be excited with amber light (Klapoetke et al., 2014). At an illumination power of 60 μW, the activity of the PV interneurons during the delivery of the high-contrast stimulus was scaled up by an average factor of 1.78 (n = 89 cells), without a significant change in time-course of sensitization (Fig. 3B and Fig. S1). To test how over-activating PV interneurons in this way impacted on the activity of PCs expressing GCaMP6f we used interleaved trials of the stimulus with and without simultaneous co-activation of ChrimsonR. The example in Fig. 3C shows a PC in which there was relatively little adaptation under normal conditions (grey bars). Increasing PV activity (purple bars) had two basic effects: the initial gain of the response was reduced, after which there was a further decrease in responsivity i.e depressing adaptation.

**Figure 3.**
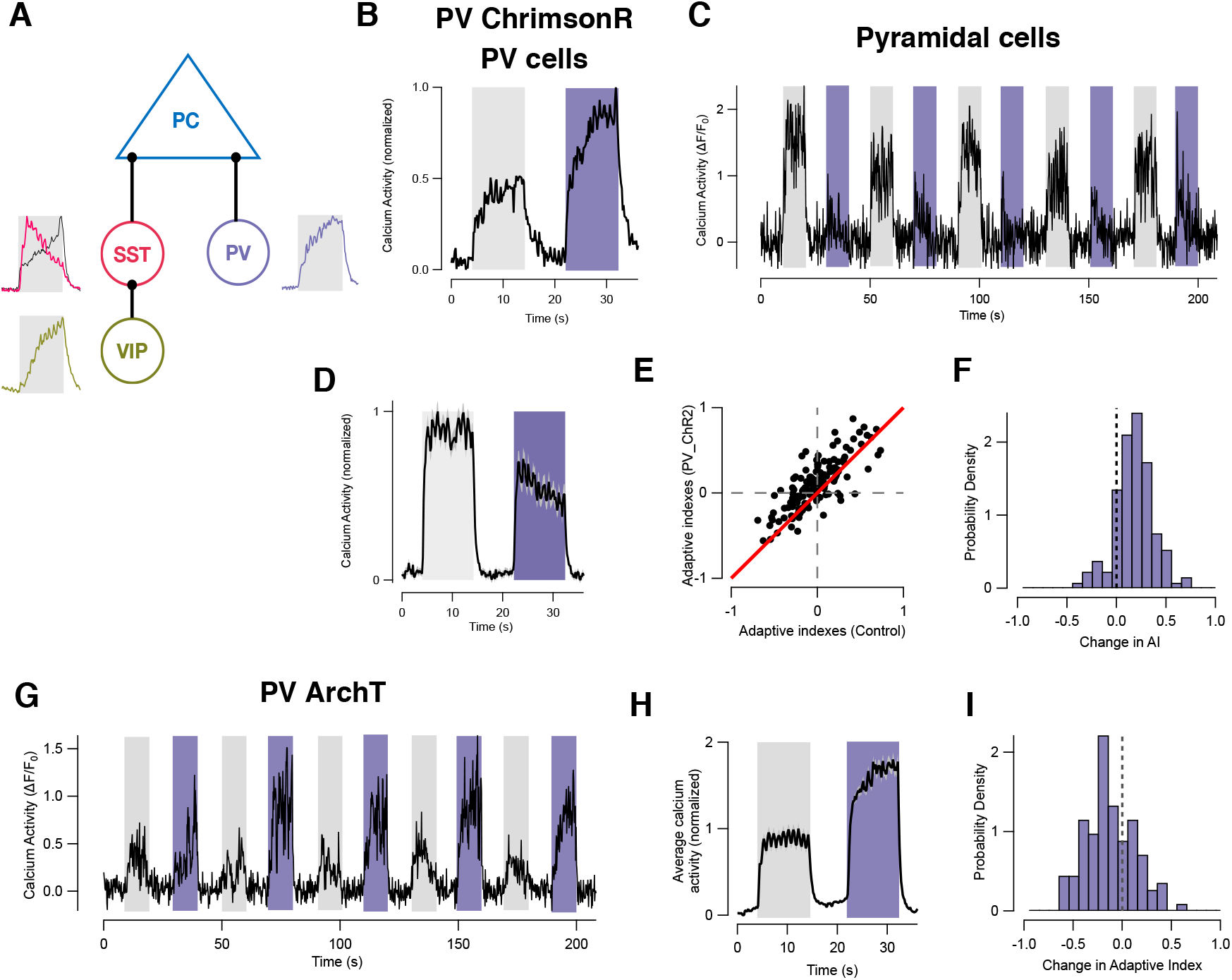
PV interneurons drive depressing adaptation in pyramidal cells. **(A)** Schematic of the major inhibitory connections in layer 2/3 of V1. The average response of each of the three class of interneuron to the high-contrast stimulus is also shown, from Fig. 2. **(B)** Population average of GCaMP6f signal in PV cells in response to the visual stimulus delivered without (*grey bar*) and with (*purple bar*) optogenetic activation using ChrimsonR (89 cells). The activity of PV cells almost doubled (see also Fig. S1). **(C)** An example of the response of a PC when optogenetically activating PV interneurons. Stimulus trials with LED illumination (purple bars) and without (grey bars) were interleaved. Note how both the amplitude and dynamics of the response were altered. **(D)** Average of paired stimulus trials from experiments shown in C (n=3 mice, 296 cells). Over-activation of PV cells reduced gain in PCs and pushed the population towards depressing adaptation. **(E)** Scatter plot showing Adaptive Index measured in PCs during activation of PV cells in relation to control. Each dot represents one neuron and the red line is unity. Most PCs demonstrate an increase in AI corresponding to depression. **(F)** Distribution of changes in AI caused by activation of PV cells expressing ChrimsonR. The distribution is shifted towards positive values (significant at p<0.009; n=3 mice, 296 cells). **(G)** An example of the response of a PC when optogenetically inhibiting PV interneurons with ArchT (purple bars). Inhibiting activity of PV cells increased gain in PCs and pushed the population towards sensitizing adaptation. **(H)** Average of paired stimulus trials from experiments shown in G (n=3 mice, 264 cells). **(I)** Distribution of changes in AI caused by inhibition of PV cells expressing ArchT. The distribution is shifted towards negative values (significant at p<0.001).

Collected results from 296 pyramidal cells (3 mice) are shown in Fig. 3D-F. Increasing PV activity reduced the average response of PCs by a factor of 1.2 ± 0.02 and caused depression to predominate (average AI = 0.12 ± 0.04; Fig. 3D, Fig. S1-4). The range of effects is shown in Fig. 3E by plotting the AI during optogenetic activation of PV interneurons against the AI measured under control conditions, each point representing one PC. Most cells fell above the line of unity (red) indicating a positive shift in AI, which was significant at p = 0.009 (t-test). From these paired measurements we calculated the change in AI caused by over-activating PV interneurons and the distribution of this change is shown in Fig. 3F: increasing the amplitude of the sensitizing response in PV interneurons pushed the large majority of pyramidal cells towards depression.

To test whether *normal* levels of activity in PV interneurons were sufficient to drive depressing adaptation in PCs we used ArchT to counteract the effect of the visual stimulus. Again, the experimental protocol used interleaved trials with and without simultaneous co-activation of ArchT (Fig. 3G). Reducing PV activity increased the average amplitude of the response in PCs (Fig. 3H, Fig. S2) and caused sensitization to predominate (Fig. 3I, Fig. S1; n = 264 cells, 3 mice, significant at p = 0.001, t-test). Together, the results in Fig. 3 demonstrate that inputs from PV interneurons modulate not only the initial gain of visual responses in PCs, but also the promote the depressing mode of adaptation.

### SST interneurons drive sensitization in pyramidal cells through two distinct mechanisms

After PV interneurons, the second major inhibitory input to pyramidal cells originates from the SST subtype, which can themselves be modulated by VIP interneurons (Pfeffer et al., 2013; Urban-Ciecko and Barth, 2016; Fig. 2F). Optogenetic activation of SST interneurons using ChrimsonR (Fig. S2A-C) had two major effects on PCs that were responsive under normal conditions: the initial gain of the response to the high-contrast stimulus was reduced, but this was followed by sensitization, as shown by the example in Fig. 4A. On average, activation of SST interneurons reduced the initial response across the population of PCs by 22% (Fig. 5B, Fig. S3 and S5) and the shift towards sensitization was significant at p = 0.0001 (Fig. 4C; t-test; n = 552 cells, Fig. S1). Suppressing SST interneurons expressing ArchT (Fig. S2D-F) had effects that were qualitatively the opposite: the initial response to the stimulus was scaled by a factor of 2.22 ± 0.10 (Fig. 4D and E), and the PC population shifted towards depression (Fig. 4F; n = 319 cells, 5 mice; significant at p = 3.3 × 10^−4^, t-test, Fig. S1).

**Figure 4.**
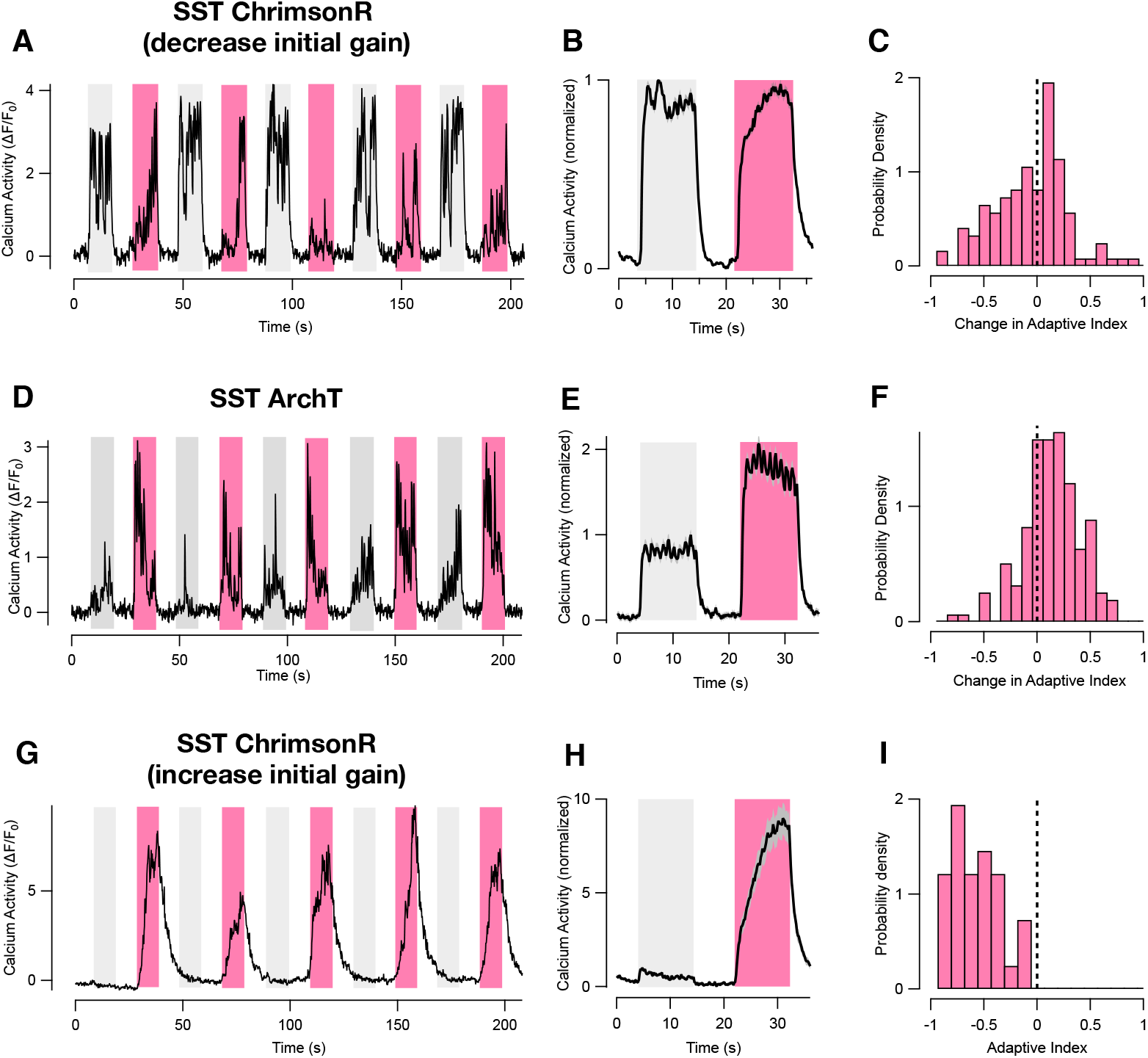
SST interneurons drive sensitization of pyramidal cells. **(A)** An example of the response of a PC when optogenetically activating SST interneurons expressing ChrimsonR. Stimulus trials with LED illumination (pink bars) and without (grey bars) were interleaved. Increasing activity of SST cells decreased the initial gain, which was followed by sensitization. **(B)** Average of paired stimulus trials from experiments shown in A (n = 3 mice, 552 cells). The variable changes in initial amplitude are shown in Fig. S6. **(C)** The distribution of changes in adaptive index of PCs during SST cell activation. The shift towards negative values of AI (sensitization) was significant at p<0.0001. **(D)** An example of the response of a PC when optogenetically inhibiting SST interneurons expressing ArchT (pink bars). **(E)** Average of paired stimulus trials from experiments shown in D (n = 3 mice, 319 cells). **(F)**The distribution of changes in AI of PCs during SST cell inhibition. The shift towards positive values of AI (depression) was significant at p = 3.34 × 10^−4^ (t-test). (**G)** An example of responses from one of the subset of PCs (12% of total) that was not normally significantly responsive to the stimulus, but became strongly responsive on activating SST interneurons expressing ChrimsonR. **(H)** Average of paired stimulus trials from experiments shown in D (n = 3 mice, 67 cells). In this subset of PCs, sensitization was correlated with a dramatic increase in gain, indicating that SST cells were exerting a disinhibitory effect. **(I)** The distribution of AI in PCs that became responsive during SST excitation through ChrimsonR: these were uniformly sensitizing (average AI = −0.393 ± 0.034).

The general picture emerging from the results in Figs. 3 and 4 is that bidirectional manipulations of activity in PV and SST interneurons push adaptive effects within the PC population from one polarity to another. This in turn suggests a model that explains variations in adaptive properties of PCs as reflecting differences in the balance between SST inputs driving sensitization and PV inputs driving depression. This model is explored further below.

A small subset of PCs (12%) sat outside this pattern. These were easily distinguished because they did not generate a significant response to the high-contrast stimulus under normal conditions but became strongly active when the stimulus was paired with activation of SST interneurons expressing ChrimsonR. These PCs were notable in sensitizing strongly (AI averaged −0.39 ± 0.03, n=67 cell; Fig. 4G-I). The key feature that distinguished this second route of sensitization was its accompaniment by a strong *increase* in initial gain (cf., Fig. 4G-I). The results in Fig. 4 therefore identify two mechanisms by which SST interneurons drove PCs to sensitize, one associated with a decrease in the initial gain (Fig. 4A-B) and the second with a dramatic increase (Fig. 4G-I). SST-expressing interneurons are diverse in terms of morphology and connectivity (Xu et al., 2013a; Tremblay et al., 2016) and a recent study has placed them into two distinct populations based on the strength of their correlations (Knoblich et al., 2019).

### VIP interneurons drive depressing adaptation in pyramidal cells

We turned next to the third major class of inhibitory interneuron expressing VIP. Direct connections between these and PCs are sparse and weak but they exert powerful indirect actions through SST interneurons (Fu et al., 2014; Niell and Stryker, 2010; Pakan et al., 2016; Dipoppa et al., 2018). This VIP->SST disinhibition pathway regulates network dynamics in V1 during changes in behavioural state, such as engagement in locomotion or arousal, reflecting the long-range inputs that VIP cells receive from other cortical areas (Niell and Stryker, 2010; Reimer et al., 2014; Vinck et al., 2015; Zhang et al., 2014).

**Figure 5.**
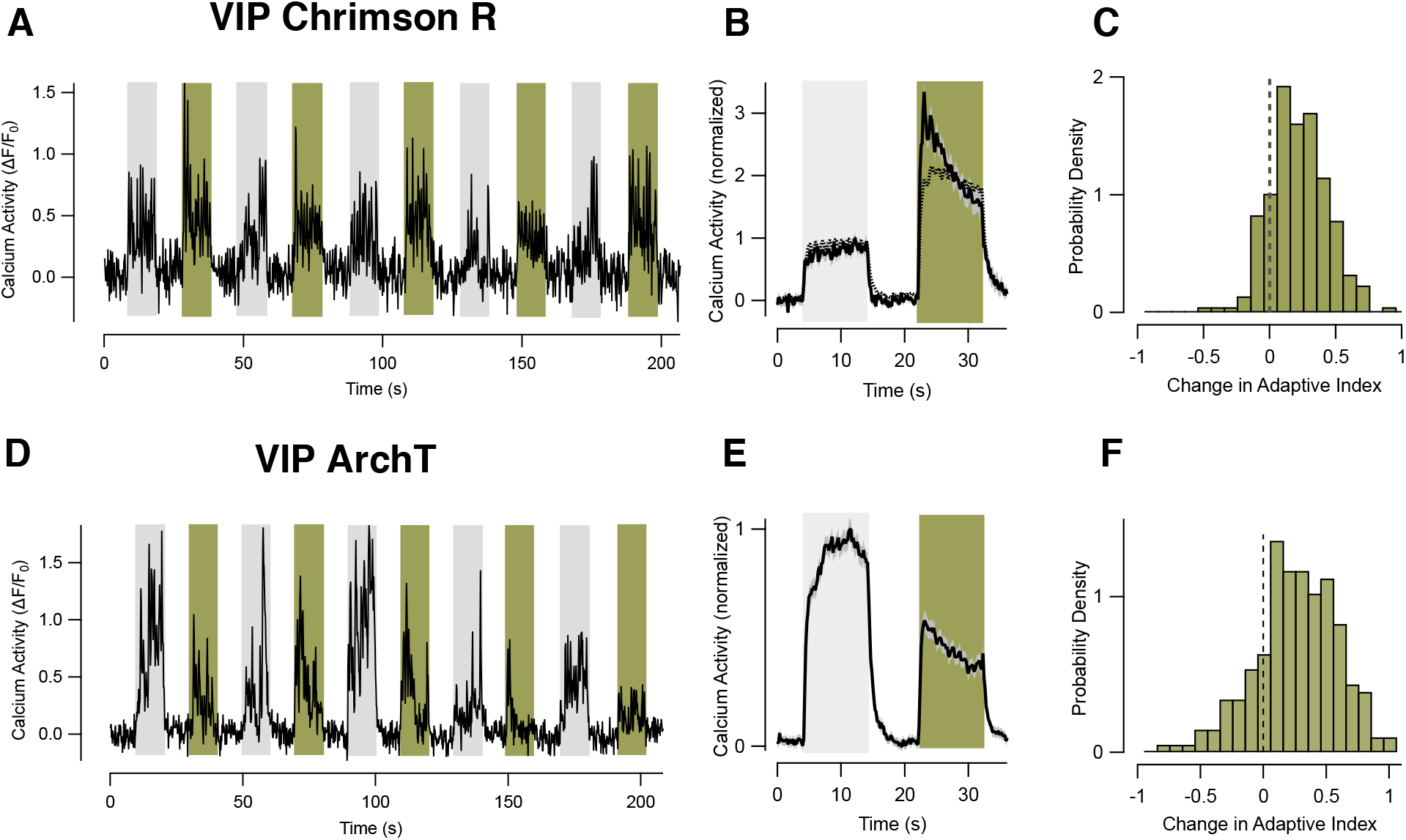
VIP interneurons drive sensitization of pyramidal cells. **(A)** An example of the response of a PC when optogenetically activating VIP interneurons expressing ChrimsonR. Stimulus trials with LED illumination (green bars) and without (grey bars) were interleaved. Increasing activity of VIP cells decreased the initial gain, demonstrating effective disinhibition, which was followed by depressing adaptation. **(B)** Average of paired stimulus trials from experiments shown in A (n = 3 mice, 414 cells). The bold trace shows the effect of 1 mW illumination power and the dashed trace 0.06 mW. **(C)** The distribution of changes in adaptive index in PCs during VIP cell activation. The shift towards positive values (sensitization) was significant at p<0.0001. **(D)** An example of the response of a PC when optogenetically inhibiting VIP interneurons expressing ArchT. **(E)** Average of paired stimulus trials from experiments shown in D (n = 3 mice, 421 cells). **(F)** Distribution of changes in AI of PCs during VIP cell inhibition The distribution was shifted towards depressing adaptation (significant at p<0.003).

To test whether the VIP->SST disinhibition circuit is also involved in adaptation, we began by manipulating the activity of VIP interneurons using ChrimsonR (Fig. 5A). When the power of the amber exciting beam was 1 mW, stimulus-driven activity in VIP interneurons expressing ChrimsonR increased by a factor of 2.37 ± 0.07 (Fig. S3A-C) and the initial response in PCs increased by 2.76 ± 0.06 (solid line in Fig. 5B, Fig. S3), consistent with strong activation of a disinhibitory pathway. The distribution of the change AI shows the population of PCs was simultaneously shifted towards depression (Fig. 5C; n = 414 cells, 3 mice; significant at p = 0.003, Fig. S1). Optogenetic excitation of VIP cells therefore had the same general effect as inhibition of SST cells (Fig. 5D-F, Fig. S1–3), consistent with a dominant role of the VIP->SST disinhibition pathway. These results are again consistent with the idea that inhibiting SST inputs into PCs shifts the balance towards PV interneurons driving depression (Figs. 3 and 4).

To test whether normal levels of activity in VIP interneurons were sufficient to exert adaptive effects in PCs, we inhibited their response to the stimulus using ArchT (Fig. 5D-F). An illumination power of 1 mW reduced the response of PCs by an average factor of 1.64 ± 0.044 (Fig. 5D, Fig. S2–3) demonstrating that this manoeuvre was effective in releasing SST interneurons from inhibition. Simultaneously, there was a shift in the distribution of AI in PCs towards depression (Fig. 5F; n = 535 cells, 3 mice; significant at p = 5 × 10^−6^, Fig. S1). This result appears paradoxical at first - over-activating or inhibiting VIP interneurons *both* shifted the population of PCs towards depression. A resolution likely lies in the observation that SST interneurons were a mixture of depressing and sensitizing (Fig. 2). VIP interneurons were uniformly sensitizing, indicating that these primarily modulate the depressing subset of SST cells. Reducing VIP activity can therefore be expected to shift the balance of adaptive effects in the SST population as a whole towards sensitization, thereby driving the population of PCs towards depression.

### Circuits determining the balance between depression and sensitization in pyramidal cells

Circuit models that integrate the observations in Figs. 1-5 are shown in Fig. 6A-C. These begin with current understanding of the most basic circuit elements in V1 (Pfeffer et al., 2013; Pi et al., 2013; Tremblay et al., 2016; Yetman et al., 2019) on which we have superimposed the network dynamics observed in normal conditions (Figs. 1 and 2) and during a variety of optogenetic manipulations (Figs. 3–5). We propose that the adaptive response in different PCs varies because of differences in the balance between direct inputs from SST and PV interneurons. (circuits 1 and 2 in Fig. 6A). PV cells are sensitizing (Fig. 2) and therefore drive depression (Fig. 3), while the majority of SST cells are depressing (Fig. 2) and drive sensitization (Fig. 4). Activity in VIP interneurons also favours depression (Fig. 5), consistent with reducing the sensitizing input of SST interneurons through the VIP->SST disinhibition pathway (circuit 3 in Fig. 6B).

**Figure 6.**
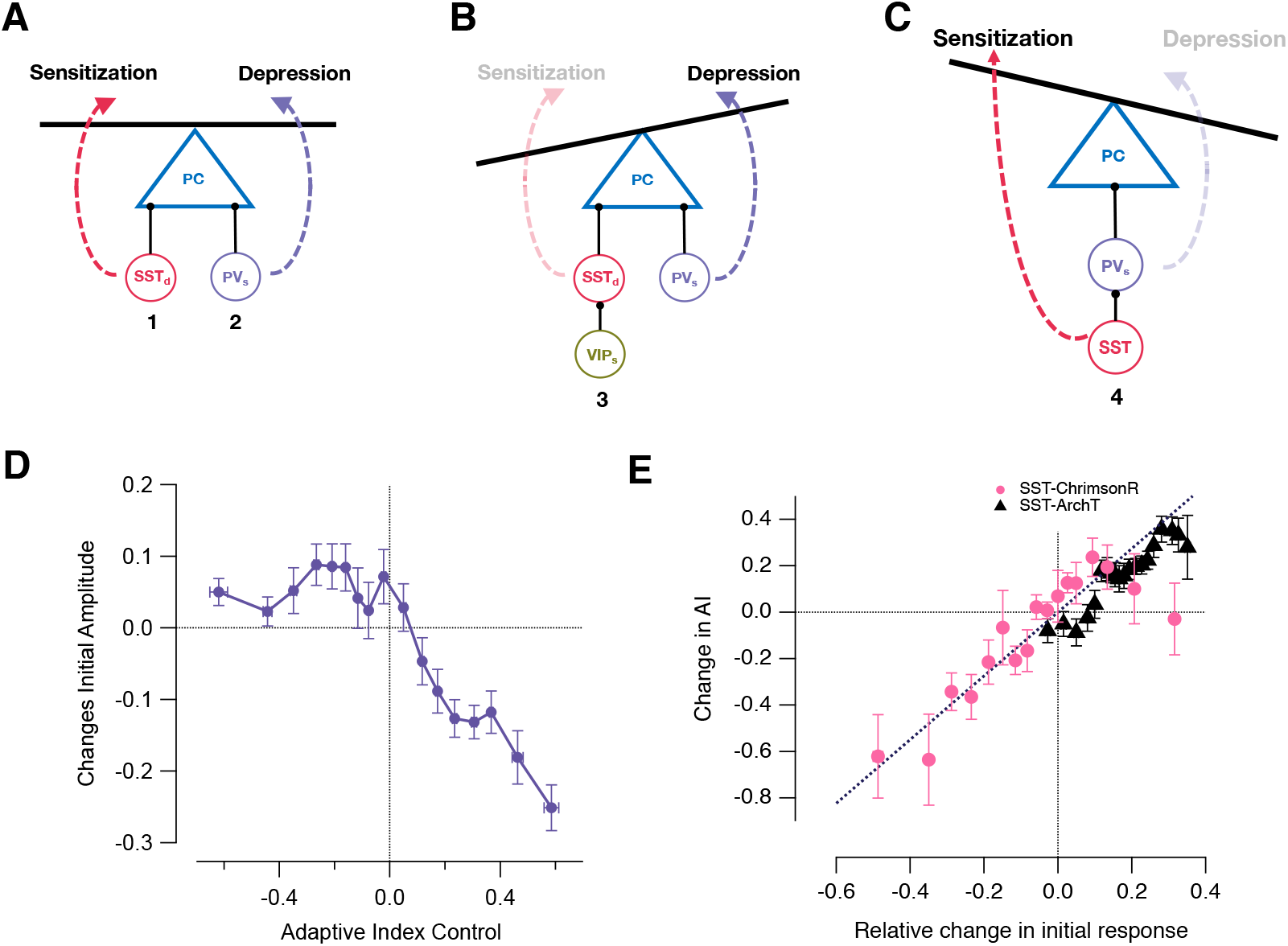
Local circuits controlling depressing and sensitizing adaptation. **(A)** We propose that the adaptive response in different PCs varies because of differences in the balance between direct inputs from SST and PV interneurons. PV_s_ cells were sensitizing and therefore drive depression. The majority of SST cells were depressing (denoted SST_d_) and drive sensitization. **(B)** Activity in VIP interneurons favoured depression, consistent with reducing the sensitizing input of SST interneurons through the VIP_s_->SST_d_ disinhibition pathway. **(C)** An SST->PV_s_ disinhibition circuit that can account for the subset of PCs (~12%) that were not normally responsive to the stimulus but then displayed a dramatic increase in gain when over-activating SST interneurons. **(D)** Plot of the relation between the change in initial amplitude of the stimulus-driven response in PCs when optogenetically activating PV interneurons (a measure of the strength of direct PV input) and the AI under normal conditions. PCs receiving the strongest PV input displayed the strongest depressing adaptation. For sensitizing PCs (AI < 0), PV activation had little effect on the initial amplitude of the response to the high-contrast stimulus, indicating that these received weak PV input. **(E)** Plot of the relation between the change in AI of PCs caused by an optogenetic manipulation and the relative change in in initial amplitude of the stimulus-driven response (a measure of the strength of direct SST input). Each point is averaged over 20-30 PCs. Pink circles show effects of activating SST interneurons using ChrimsonR: the stronger the SST input the stronger the sensitization (Correlation coefficient, r = 0.85). Black triangles show effects of inhibiting SST interneurons using ArchT. The greater the increase in gain, the stronger the shift to the depressing mode of adaptation (Correlation coefficient, r = 0.93). The dashed line is a fit constrained to pass through the origin. These results support the prediction of the model in A, that PCs with the strongest *direct* input from SST cells will also have the strongest tendency to sensitize.

A key prediction of the model in Fig. 6A is that PCs with the strongest tendency to depress also receive the strongest input from PV cells. We tested this idea by separating PCs according to AI and then assessing how efficiently they were inhibited by optogenetic activation of PV cells expressing ChrimsonR. Fig. 6D demonstrates that, as predicted by the model, the more strongly a pyramidal cell depressed (AI > 0), the stronger the inhibitory input it received from PV interneurons. The initial gain of sensitizing PCs (AI < 0) was far less sensitive to PV activation, indicating that these neurons received weak PV input.

A second prediction of the “balance” model in Fig. 6A is that PCs with the strongest *direct* input from SST cells will have the strongest tendency to sensitize, and this was confirmed by the analysis in Fig. 6E. Here we grouped PCs according to the change in gain caused by optogenetic activation of SST interneurons, assessed from the initial amplitude of the response to the high-contrast stimulus. The change in AI for these different groups is plotted as the pink circles and it can be seen that the stronger the SST input the stronger the sensitization (Correlation coefficient, r = 0.85). The same analysis was carried out when SST interneurons were inhibited through ArchT (black triangles in Fig. 6E). The greater the increase in gain, the stronger the shift to the depressing mode of adaptation (Correlation coefficient, r = 0.93), indicating that SST inputs were normally driving sensitization. These results provide further evidence for the idea that the relative strength of SST and PV inputs determine whether there is an increase or decrease in gain during the presentation of the high-contrast stimulus, at least in most PCs.

The balance model does not account for the subset of PCs (~12%) that were initially silent but then displayed a dramatic *increase* in gain when over-activating SST interneurons (Fig. 4G-I). Clearly, these PCs do not receive significant *direct* input from SST cells, which must instead act in a disinhibitory mode. This is very likely to occur through PV cells (Fig. 6C) given that these provide strong inhibitory inputs to all PCs (Pfeffer et al., 2013; Yetman et al., 2019) and an SST->PV disinhibition pathway has been shown to regulate visual responses in some PCs in V1 (Cottam et al., 2013; Pfeffer et al., 2013; Rikhye et al., 2017). SST interneurons can therefore drive sensitization in at least two distinct groups of PCs, one associated with a decrease in initial gain and the second with an increase (Fig. 4).

### Locomotor behaviour increases gain while maintaining the balance of adaptive effects across the population of pyramidal cells

Changes in the gain of PC responses are determined not only by the history of the external stimulus, but also by changes in the internal state of the animal. Most notably, dramatic increases gain occur when an animal transitions from a “resting” or disengaged state to an “active” or aroused state signaled by locomotion or an increase in pupil diameter (Niell and Stryker, 2010; Vinck et al., 2015; Pakan et al., 2016). These changes in cortical processing reflect long-range inputs such as those from the basal forebrain and locus coeruleus (reviewed in Ferguson and Cardin, 2020). How do changes in internal state alter the properties of adaptation across the population of PCs? To investigate this question we compared visual responses measured at rest and during locomotion, categorized according to a threshold velocity (see Methods). Fig. 7A shows the activity of a PC in which every second stimulus was applied with inhibition of SST interneurons through ArchT (pink bars). Note that the gain of control responses was increased during locomotion (grey bars), although not to the extent caused by the optogenetic manipulation.

**Figure 7.**
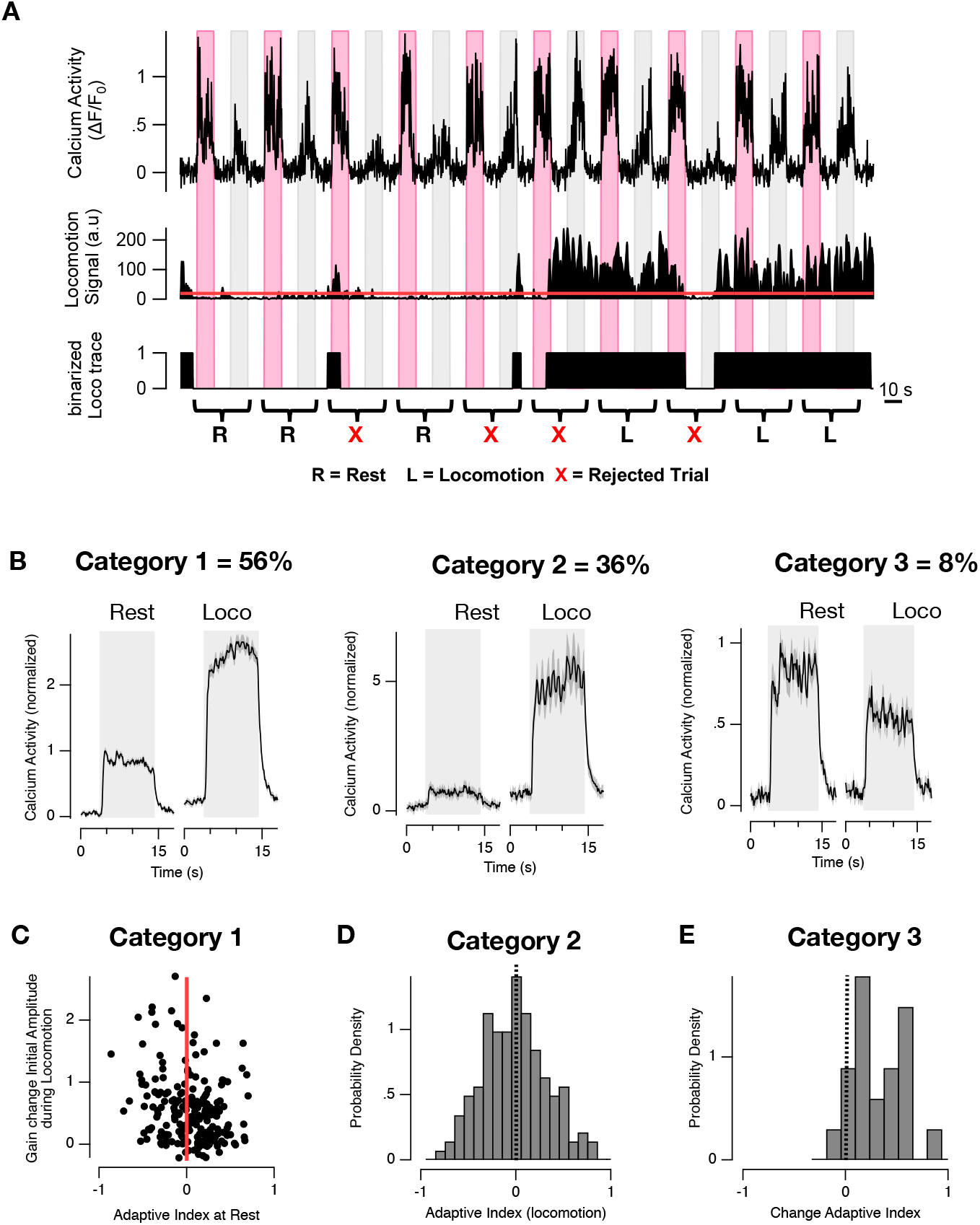
Locomotion activates both depressing and sensitizing pyramidal cells. **(A)** An example of the responses from a PC in a mouse expressing ArchT in in SST interneurons (top trace). Control trials (grey bars) were interleaved with trials with illumination of ArchT (pink bars). Locomotion was intermittent (middle trace) and was binarized (bottom) by setting a threshold (red line). A pair of trials was counted as occurring at rest (R, 32% of total) if there was no coincident locomotion and only counted as locomotion (L, 42%) if this was continuous during both presentations of the stimulus. A pair was rejected for analysis (X, 26%) if there was any coincident switch between R and L, and any imaging session with less than 3 trials for both rest and locomotion conditions was excluded from this dataset. **(B)** Averaged responses at rest and during locomotion within three category of PC, collected from 5 mice. Category 1 - responsive to stimulus at rest with gain increase during locomotion (n = 253); category 2 - not responsive to stimulus at rest with gain increase during locomotion (n = 162); category 3 - response to stimulus but inhibited during locomotion (n = 34). Note that there was no net time-dependent change in activity across the population of PCs, indicating that depression and sensitization were balanced. **(C).** The change in initial amplitude of the response during locomotion plotted as a function of AI at rest for PCs in category 1. Both depressing and sensitizing neurons experienced increases in gain during locomotion. **(D)** The distribution of AIs for PCs in category 2 during locomotion. The mean was not significantly different from zero (AI = −0.042 ±.028; n = 162 cells; t-test). **(E)**The distribution of the change in AI for PCs in category 3 (mean = 0.22 ± 0.06; n = 34) showing a significant shift towards depression (p = 0.006; t-test).

All the PCs that were responsive to the high-contrast stimulus, either at rest or during locomotion, could be placed into one of three categories (Fig. 7B); category 1 (56%) were responsive at rest and showed a significant increase in gain; category 2 (36%) did not show a significant response to the stimulus at rest but became responsive during locomotion, and category 3 (8%) were inhibited during locomotion. Cells in category 2 might also be thought of as one extreme of category 1, but with gain at rest so low as to make responses difficult to detect without averaging. For PCs in category 1, the gain increase during locomotion increased the initial amplitude of the response by a factor 2.49 ± 0.18. Crucially, this reflected an increase in gain of both depressing and sensitizing neurons PCs (Fig. 7C). For PCs in category 2, the distribution of AI during locomotion was also balanced between depression and sensitization (Fig. 7D).

These observations are not easy to reconcile with the idea that locomotion acts solely through the VIP->SST disinhibition pathway, since this drove depressing adaptation (Fig. 5A and 6B). Rather it seems that locomotion also increases the gain of some PCs through a circuit that drives sensitization. The best candidate for this is the SST->PV disinhibition pathway uncovered by activating SST interneurons using ChrimsonR (Fig. 4G-I; Fig. 6C; Fig. 7A). This conclusion is also consistent with two observations made recently by Dipoppa et al. (2018): locomotion induces strong activation in almost as many SST interneurons as VIP cells (58% vs 70%) and as many PV neurons are inhibited as activated (see their Fig. 2). An increase in gain acting across both depressing and sensitizing neurons was also observed in PCs that only became responsive during locomotion (Fig. 7D), indicating that these are also a mixed population in which either of the two disinhibition pathways predominates.

Finally, the small subset of PCs experiencing a *decrease* in gain during locomotion were distinctive in that almost all were pushed towards depression (Fig. 7E). This might reflect PCs in which the dominant inhibitory input is from PV neurons *activated* during locomotion, given that these were almost uniformly sensitizing (Fig. 2A) and their optogenetic activation also reduced gain while driving depression (Fig. 3B-F).

## Discussion

This investigation demonstrates that pyramidal cells in layer 2/3 of V1 adapt to a high-contrast stimulus to varying degrees and with opposing forms of plasticity. While many PCs initially respond with high gain followed by gradual depression over periods of several seconds, these are balanced by cells that respond with low gain but then sensitize (Fig. 1). These different modes of adaptation reflect the dynamics of signals in three classes of interneuron (Fig. 2), and a variety of optogenetic manipulations (Figs. 3–5) were consistent with a model in which the net adaptive effect reflects the balance between direct inputs from PV interneurons, driving depression, and a subset of SST interneurons, driving sensitization (Fig. 6). A second level of control occurs indirectly, through disinhibitory circuits, the VIP->SST pathway driving depression (Fig. 5) and the SST->PV pathway driving sensitization (Fig. 4).

The change in internal state associated with the onset of locomotion increases the gain of PCs expressing both major forms of plasticity, depression and sensitization (Fig. 7). We propose, therefore, that long-range inputs active during locomotion drive both disinhibitory circuits, as schematized in Fig. 8. This model provides an integrated view of the mechanisms underlying changes in gain in response to changes in cortical state and during opposing forms of adaptation, at least for the ~92% of PCs in which responsivity is enhanced during locomotion. The general picture is that distinct subsets of PCs are purposed for signalling increases or decreases in contrast, but all are engaged during locomotion to maintain the balance between opposing forms of adaptation (Figs. 8) and relatively constant activity averaged across the population of PCs (Fig. 1 and Fig. 7).

**Figure 8.**
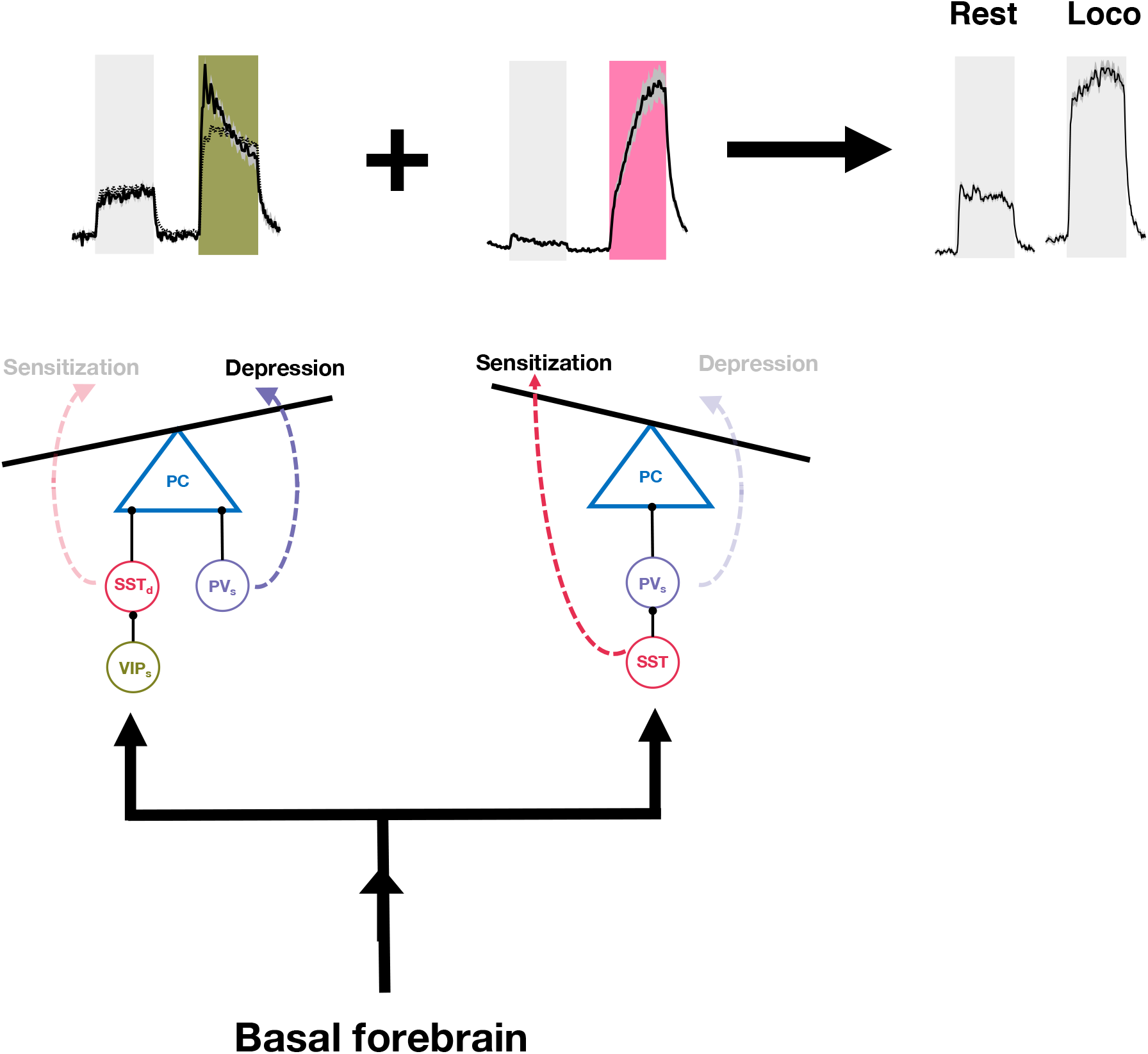
Interactions between the inhibitory microcircuits controlling opposing forms of adaptation to an external stimulus and changes in behavioural state. The transition from rest to a locomotor state increases the gain of visual responses in both depressing and sensitizing PCs (top left), so that the average activity over the population is relatively constant (top right). Circuits based on the results in this paper can account for these two forms of plasticity. We propose that long-range inputs from the basal forebrain that are active during locomotion drive two disinhibitory circuits: the VIP->SST circuit increases gain while driving some PCs towards depressing adaptation, while the SST->PV circuit increases gain while driving PCs to sensitize. The upper panels schematize this model by analogy with the observed effects of optogenetic activation of VIP interneurons (top left, green shading, from Fig. 5) and SST interneurons (top middle, pink shading, from Fig. 4G-I).

### Circuit mechanisms of depressing and sensitizing adaptation

It is only recently that sensitization has been identified as an important adaptive mechanism (Solomon and Kohn, 2014) and the neural circuit in which it has been characterized best is the retina (Kastner and Baccus, 2011; Nikolaev et al., 2013; Appleby and Manookin, 2019). Both depressing and sensitizing adaptation begin in the synaptic compartments of bipolar cells that use glutamate to transmit the visual signal to the retinal output neurons, the ganglion cells. These synapses are a key site of gain control in the retina and receive direct input from inhibitory amacrine cells (Johnston et al., 2019). The major cause of contrast adaptation as a decrease in synaptic gain is depletion of vesicles available for release (Burrone and Lagnado, 2000; Nikolaev et al., 2013b; Ozuysal and Baccus, 2012), while sensitization reflects presynaptic depression in inhibitory amacrine cells providing fast (GABAergic) feedback (Appleby and Manookin, 2019; Kastner et al., 2019; Nikolaev et al., 2013). The balance between these two effects varies according to the frequency of the stimulus and the inhibitory microcircuit in which the bipolar cell terminal is embedded, leading to partial segregation of depressing and sensitizing signals transmitted to RGCs ( Nikolaev et al., 2013; Kastner and Baccus, 2013). Here we have concentrated on the dynamics of inhibitory interneurons, but might modulation of excitatory inputs also contribute to contrast adaptation in V1?

Excitatory synaptic inputs from thalamocortical neurons to PCs are known to be subject to presynaptic depression although the time-scales are of the order of hundreds of milliseconds rather than the several seconds over which depressing adaptation develops in V1 (Abbott, 1997; Cimenser and Miller, 2014; Varela et al., 1999). A recent study has found evidence of presynaptic GABA_A_ receptors that modulate thalamocortical inputs in Layer 4 of rat V1 (Wang et al., 2019), but there is no evidence of direct inhibitory connections to excitatory boutons so these are likely to be activated indirectly by GABA spillover. A more complete picture of the mechanisms underlying increases and decreases in the gain of PCs will require better understanding of the role of excitatory inputs during adaptation and this might come with *in vivo* imaging of glutamate release using iGluSnFR (Marvin et al., 2013) or presynaptic calcium signals using SyGCaMPs (Dreosti et al., 2009; Schröder et al., 2019). It will be just as important to understand the cellular processes controlling adaptive responses in the different types of interneuron.

### What is the function of sensitization?

By operating simultaneously in different neurons, depressing and sensitizing forms of adaptation improve the overall rate of information transfer through the optic nerve when the input contains fluctuating contrast, as it does in the real world (Kastner and Baccus, 2011). This is because the subset of neurons that respond to an increase in contrast with low initial gain but then sensitize allow the retina to signal future *decreases* in contrast more effectively than neurons that depress. It may be that opposing forms of plasticity in V1 also provide a general strategy for “hedging bets” in the face of an unpredictable future. A second possibility is that different modes of adaptation play more specialized roles. In the retina of primates, for instance, sensitization is predominantly a property of midget ganglion cells in the fovea, allowing them to encode natural movies more accurately than depressing ganglion cells with larger receptive fields (Appleby and Manookin, 2019). Similarly, it will be interesting to explore how the functional properties of sensitizing and depressing PCs in V1 might differ and the roles that each might play in encoding a wider range of stimuli.

Here we have proposed circuit models that explain a range of experimental observations (Fig. 6 and 8), but only under limited stimulus conditions. The activity of different types of interneuron is differentially affected by changes in stimulus size and by different behavioural states, which is in turn likely to alter the adaptive effects expressed in PCs. For instance, larger stimuli (>30°) increase the activity of SST interneurons particularly strongly (Dipoppa et al., 2018; Pakan et al., 2016). Do large stimuli drive the PC population towards sensitization (Fig. 6)? Or might correcting mechanisms continue to maintain the balance between depressing and sensitizing adaptation, as observed using a 20° stimulus (Fig. 1)? Similarly, how do the effects of locomotion compare with other changes in behavioural or cognitive state, such as those associated with spatial attention or conditioning to a reward stimulus? A recent study indicates that sensitization becomes the dominant form of adaptation when a mouse is conditioned to a rewarded stimulus (Keller et al., 2017). The results we have presented may provide a framework for understanding the circuit mechanisms underlying this effect and perhaps other changes in internal state that alter the processing of stimuli originating from the external world.

## Methods

### Experimental Model and Subject Details

All experimental procedures were conducted according to the UK Animals Scientific Procedures Act (1986). Experiments were performed at University of Sussex under personal and project licenses granted by the Home Office following review by the Animal Welfare and Ethics Review Board (AWERB).

Experiments were performed on 24 adult transgenic mice of either sex (4 – 10 months) expressing the Cre recombinase in specific subsets of interneuron on a C57BL/6J background. Results are reported from 8 VIP-Cre mice (VIP tm1(cre)Zjh/J Jackson #), 7 PV-Cre (Pvalb tm1(cre)Arbr/J, Jackson #008069) and 9 SST-Cre (SST tm2.1(cre)Zjh/J, Jackson #013044). Mice were housed individually on inverted light-dark cycles and had access to a complex enriched environment after the end of each imaging session. This environment was large (~80 × 40 × 40 cm) and contained a number of toys, platforms and tubes that encouraged motor activity. As a result, mice were engaged in running the large majority of the time during an imaging experiment.

### Animal Preparation and Virus Injection

We prepared mice for multiphoton imaging of the visual cortex following established protocols (Goldey et al, 2014). Surgeries were performed on adult mice (P60-P90) in a stereotaxic frame under isoflurane anaesthesia (induction at 4% and 1-2% during surgery). A titanium head plate was attached to the skull, followed by a 3 mm craniotomy and durotomy to expose the brain. We then injected different combinations of viruses for imaging and manipulating cellular activity. To image calcium activity in pyramidal cells we used AAV1.CaMKII.GCaMP6f.WPRE.SV40 (n = 12 mice, titer 4.10^11^ GC/ml) or AAV5.CaMKII.GCaMP6f.WPRE.SV40 (n = 6 mice, titer 4.10^11^ GC/ml) viruses. To image calcium activity in interneurons we expressed AAV9.CAG.Flex.GCaMP6f.WPRE.SV40 in Cre lines (PV-Cre, VIP-Cre, SST-Cre, n = 6 mice, titer 2.10^12^ GC/ml). To excite interneurons optogenatically we used rAAV9/Syn.Flex.ChrimsonR.tdTomato (n = 12 mice, 2.10^12^ GC/ml) and to inhibit we used AAV5.CBA.Flex.ArchT-tdTomato.WPRE.SV40 (n = 12 mice, titer 2.10^12^ GC/ml). Viruses were injected with a beveled micropipette attached to a stereotaxic micromanipulator. Injections were performed at a single site in monocular V1 (2.4-2.8 mm lateral and 3.5-4.0 mm posterior from Bregma) and at 250-350 μm depth. A volume of 1μl of virus was injected at a single site, at a speed of 20-50 nl/min and the micropipette was retracted after a 15-20 min waiting period. A cranial window assembly was made of 2 round coverslips, (3 mm diameter, thickness #1) bonded to each other and then to a 5 mm round coverslip, (thickness #1) using optical glue (ThorLabs NOA63). The window was then placed in the craniotomy and sealed with Vetbond and dental cement. Mice were returned to their home cage for 2 weeks following surgery. At least one week before imaging started mice were habituated to head fixation under the microscope, as well as the spherical treadmill. The two LED monitors were turned on (grey screen and occasionally visual stimuli). Imaging of neural activity began 3-5 weeks after surgery and injection.

### Muiltiphoton imaging *in vivo*

Fluorescence was measured with a two-photon microscope (Scientifica SP1, galvonometer mirrors) controlled using Scanimage 5 software (Vidrio Technologies) and using a Nikon 16 × water-immersion objective (0.8 NA). The framerate was set at 6.07 Hz in bidirectional mode, for an image resolution of 256 by 200 pixels and data acquired with a 250 MHz digitizer (National Instruments). The light source was a Ti:sapphire laser (Chamelon 2, Coherent), tuned to 940 nm. Laser power under the objective never exceeded 70 mW. Imaging was carried out at a depth of 150-300 μm below the surface of the brain.

### Visual stimuli

Visual stimuli were generated using the python library PsychoPy (Peirce, 2007) running on Linux and displayed on two LED backlit monitors (BenQ XL2410T, iso-luminant at 25 cd/m^2^, 120Hz refresh rate, gamma corrected) each positioned 14.5 cm from the eyes and a 45° angle from the longitudinal axis of the animal. For all experiments, stimuli were sinusoidal gratings drifting upwards (80% contrast, 10 s duration). The size of the stimulus was set at 20 degrees of visual field, and spatial and temporal frequencies were fixed at 0.04 cpd and 1 Hz respectively. This stimulus parameters have been shown to engage all interneurons in V1 (diPoppa et al., 2018). The standard experimental protocol consisted of 10 presentations of the stimulus with a 10 s interstimulus interval consisting of a uniform grey screen of the same mean luminance. When testing an optogenetic manipulation 20 presentations of the stimulus were made, every second paired with illumination through the amber LED.

### Monitoring of Locomotion

Mice were head-restrained but free to run on a spherical air-supported treadmill (Dombeck et al., 2007). The speed of locomotion was measured with an optical mouse (GT650, gaming mouse) positioned in front of the treadmill. Signals from the mouse were conditioned through an Arduino and digitized in parallel with the signals from the detectors of the microscope. For each experiment, post-hoc analysis of the locomotion trace from the optical reader allowed us to binarized it into “still” and “moving”. First, the trace was smoothed and the baseline calculated as the mode of all the bins, which corresponded to periods of rest. The trace was then averaged over bin periods of 2.5 s. A bin was classified as “moving” if its value was greater than a threshold of three standard deviations above the baseline. To compare visual responses in the “resting” and “moving” states we only used stimulus epochs in which the mouse was continuously in that state and for at least three trials per session. (i.e responses were not used for analysis if the mouse started or stopped moving during the trial). An example of how these criteria were applied is shown in Fig. 7A. The method of housing mice (described above) encouraged motor activity and they were capable of running consistently during our imaging sessions.

### Analysis of two-photon calcium imaging data

Raw movies acquired as TIFF files were registered and segmented into Regions of Interest (ROIs) using the Suite-2P package running in MATLAB 2015b and then analyzed further using custom written code in Igor Pro 8 (Wavemetrics) including the analysis package SARFIA (Dorostkar et al., 2010). For each cell, the average fluorescence signal within the ROI, F, was background-corrected by averaging the signal in a “halo” of pixels extending ~1.5 times the width of the ROI, excluding any that fell within another ROI. The relative intensity of this background signal was then reduced by a ‘contamination ratio’ equivalent to the space occupied by the ROI itself, estimated at 0.5 here and elsewhere (Kerlin et al., 2010; Peron et al., 2015), after which it was subtracted from the raw signal. Activity traces were expressed as relative changes in fluorescence by dividing the change in fluorescence at each time point by the baseline fluorescence (ΔF/F_0_). The baseline F_0_ was usually computed as the mode of the entire activity trace when this was close to the minimum of that trace.

For each cell, the Pearson’s correlation coefficients was computed between it’s activity trace (ΔF/F_0_) and a stimulus trace binarized as either stimulus off (0) or stimulus on (1). In order to determine if the computed correlation was statistically significant, a bootstrap technique was employed in which the stimulus traces were circularly shifted from a random origin before recomputing correlation with activity as described before (Dipoppa et al., 2018). This process was repeated 1000 times, from which a probability distribution of correlation coefficients was obtained, allowing a p-value to be calculated for the coefficient value from the actual trace. The threshold for significant positive or negative correlation was set at p<0.05. Measurements of adaptive properties were confined to cells which were significantly positively correlated with the stimulus.

The Adaptive Index (AI) was calculated for each 10 s stimulus trial as the normalized ratio between the average responses during the first 2 cycles and the last 2 cycles. Measurements were rejected if there was no significant response during the trial. Therefore, cells for which optogenetic manipulation supressed responses completely were excluded from analysis. The average AI for each cell under any given condition (with or without locomotion or an optogenetic manipulation) was calculated as the average across multiple stimulus trials, usually at least 10. Changes in the initial gain were calculated from the average response during the first 2 cycles of the 1 Hz stimulus.

### Optogenetics

Red-shifted optogenetic activators (ChrimsonR or ArchT) were excited through the objective using an amber LED (Thorlabs, 590 nm, M590l3) controlled through a high-power LED driver (DC2200). To prevent contamination of GCaMP signals, the LED was pulsed to deliver light only during the turning phase of the x-mirror of the microscope, as monitored through the position signal of the mirror controller (Cambridge Technology). An Arduino read the position signal from the mirror controller (Cambridge Technology) and delivered a TTL signal for the LED driver. Under our usual imaging conditions, the LED power was was pulsed at 2 KHz, each pulse lasting ~0.1 ms. Where the effect of the optogenetic manipulation was to inhibit PCs, it was important not to completely suppress the initial response so as to quantify a change in AI. LED power was therefore calibrated during each imaging session to ensure the initial response of most PCs to the visual stimulus was not reduced by more than ~50%. This was achieved using powers of 10μW-1mW.

### Contact for Reagent and Resource Sharing

Further information and requests for resources and reagents, including custom-written code for data analysis, are available upon request from the corresponding author. Enquiries should be directed to (and will be fulfilled by) the Lead Contact, Leon Lagnado.

## Supplementary Information

**Figure S1.**
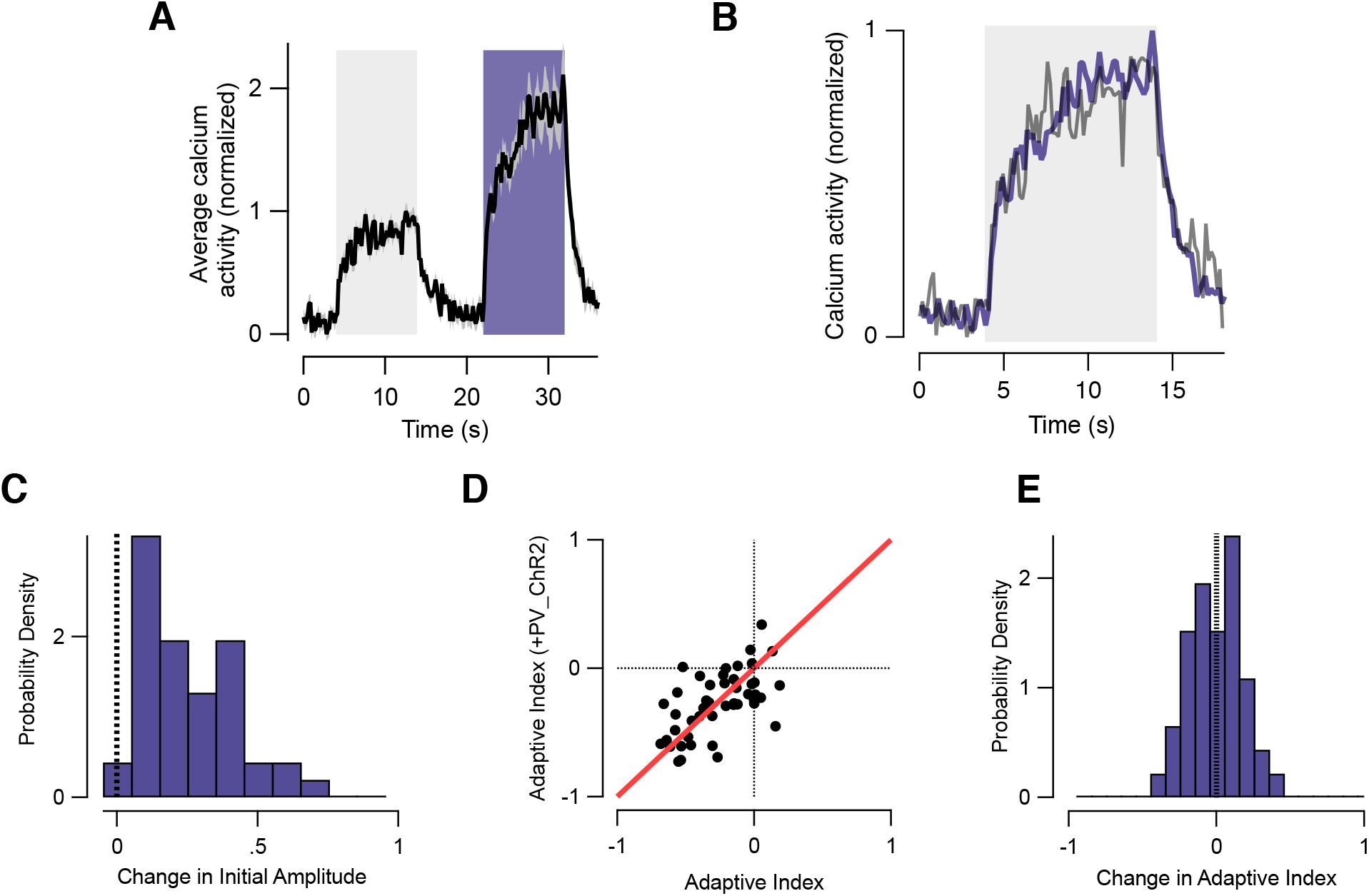
PV activation with ChrimsonR increases activity without changing the kinetic of adaptive changes. **(A)** Population trial-average of n = 89 PV interneurons responses to visual stimulus (left) and during activation of PV interneurons with ChrimsonR (right, purple bar). **(B)** The response during PV activation from A scaled down by a factor 0f 1.78 (purple) superimposes closely on the control response (grey). **(C)** Probability density distribution of relative changes in initial Amplitude of PVs during their activation with ChrimsonR. Note that initial amplitude was significantly increased by a factor of 1.84 ± 0.27 during ChrimsonR activation (n = 89 cells, p = 1.64 × 10^−10^, t-test) **(D)** Scattered plot of Adaptive index measured from the population in panel A (abscissa represent AI in control condition and ordinate during optogenetic activation of PV. Negative AI values represent sensitizing-adaptation while positive AI values represent depressing-adaptation). Each black dot represents an individual neuron and the red thick line corresponds to unity. **(E)** Plot of the probability density distribution of relative changes in adaptive index of PVs during activation with ChrimsonR. Note that activation of PVs with ChrimsonR did not significantly affect the adaptive index of individual interneurons although both their initial and average response increased (mean AI = −0.05 ± 0.03, p= 0.054, n=89, t-test)

**Figure S2.**
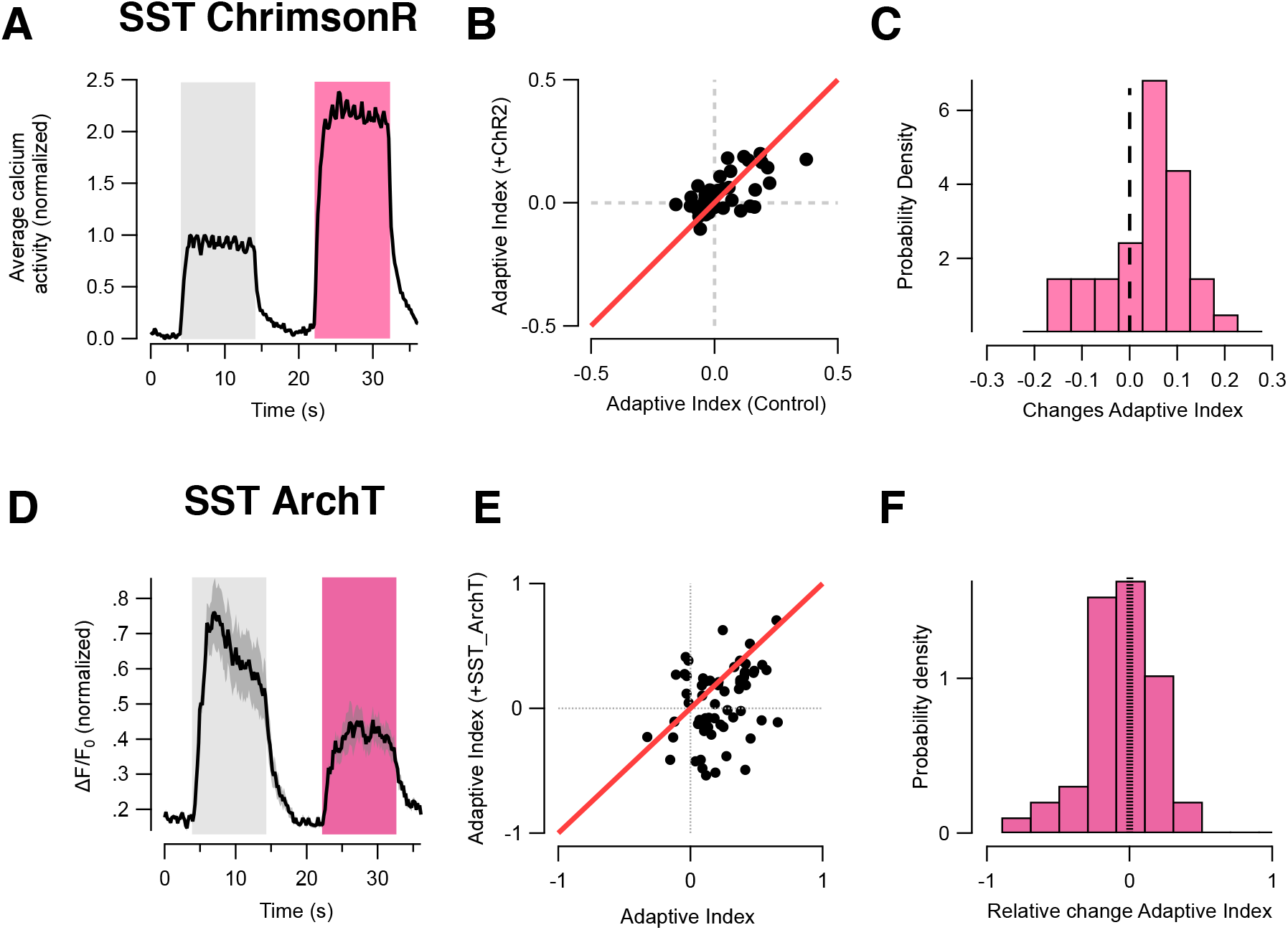
Somatic activity of SST interneurons during optogenetic activation and inhibition. **(A)** Population trial-average of n = 68 SST interneurons responses to visual stimulus (left) and during activation of SST interneurons with ChrimsonR (right, pink bar). **(B)** Scattered plot of Adaptive index measured from the population in panel A (abscissa represent AI in control condition and ordinate during optogenetic activation of SST. Negative AI values represent sensitizing-adaptation while positive AI values represent depressing-adaptation). Each black dot represents an individual neuron and the red thick line corresponds to unity. **(C)** Probability density distribution of relative changes in adaptive index of SSTs during activation with ChrimsonR. Activation of SSTs with ChrimsonR shifted the adaptive index of most interneurons towards depression depression (mean AI = 0.142 ± 0.013 p = 3.6 × 10^−4^, n=68, t-test). (**D-F**) Similar to panel A-C but using ArchT instead of ChrimsonR to inhibit SST activity. Inhibition of SST with ArchT shifted the population of SSTs towards sensitizing-adaptation (mean AI = −0.175 ± 0.034, p= 6.65 × 10^−6^, n=84, t-test).

**Figure S3.**
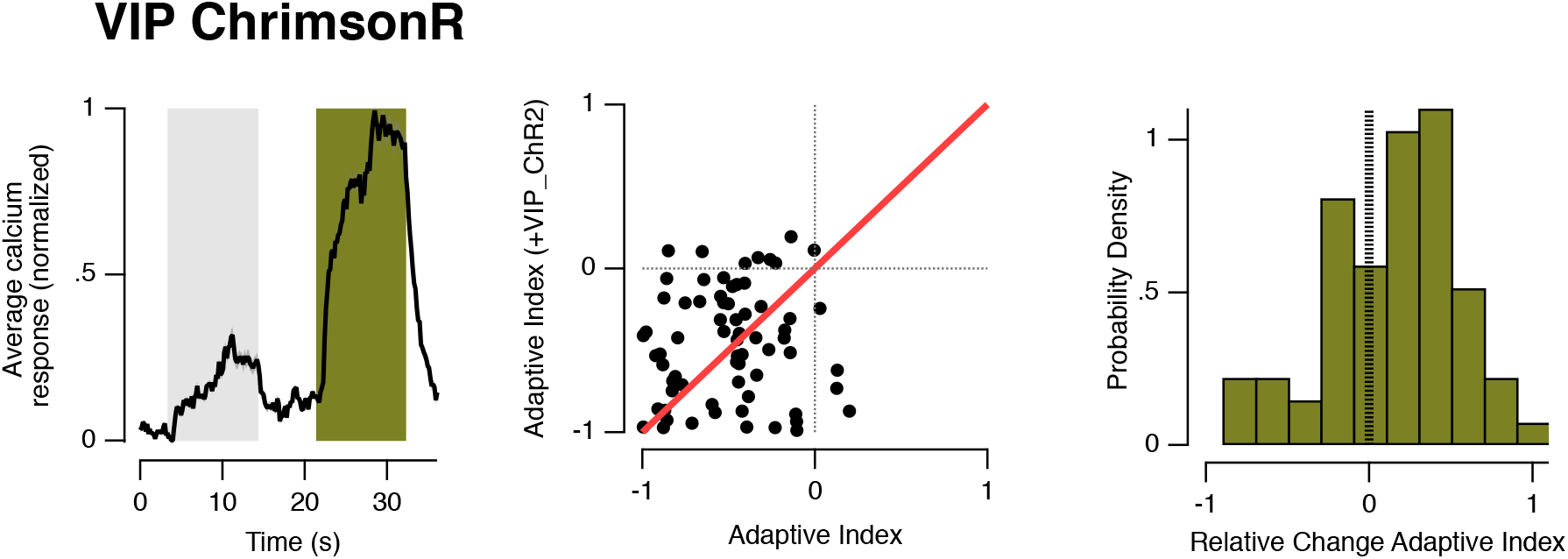
VIP activation with ChrimsonR increases activity without changing the kinetic of adaptive changes. **(A)** Population trial-average of n = 96 VIP interneurons responses to visual stimulus (*left*) and during activation of VIP interneurons with ChrimsonR (*right, green bar*) **(C)** Scattered plot of Adaptive index measured from the population in panel A (abscissa represent AI in control condition and ordinate during optogenetic activation of VIP. Negative AI values represent sensitizing-adaptation while positive AI values represent depressing-adaptation). Each black dot represents an individual neuron and the red thick line corresponds to unity. **(D)** Plot of the probability density distribution of relative changes in adaptive index of VIPs during activation with ChrimsonR. Note that activation of VIPs with ChrimsonR did not significantly affect the adaptive index of individual interneurons although both their initial and average response increased (mean AI = 0.033 ± 0.053, p= 0.535, n=96, t-test)).

**Figure S4.**
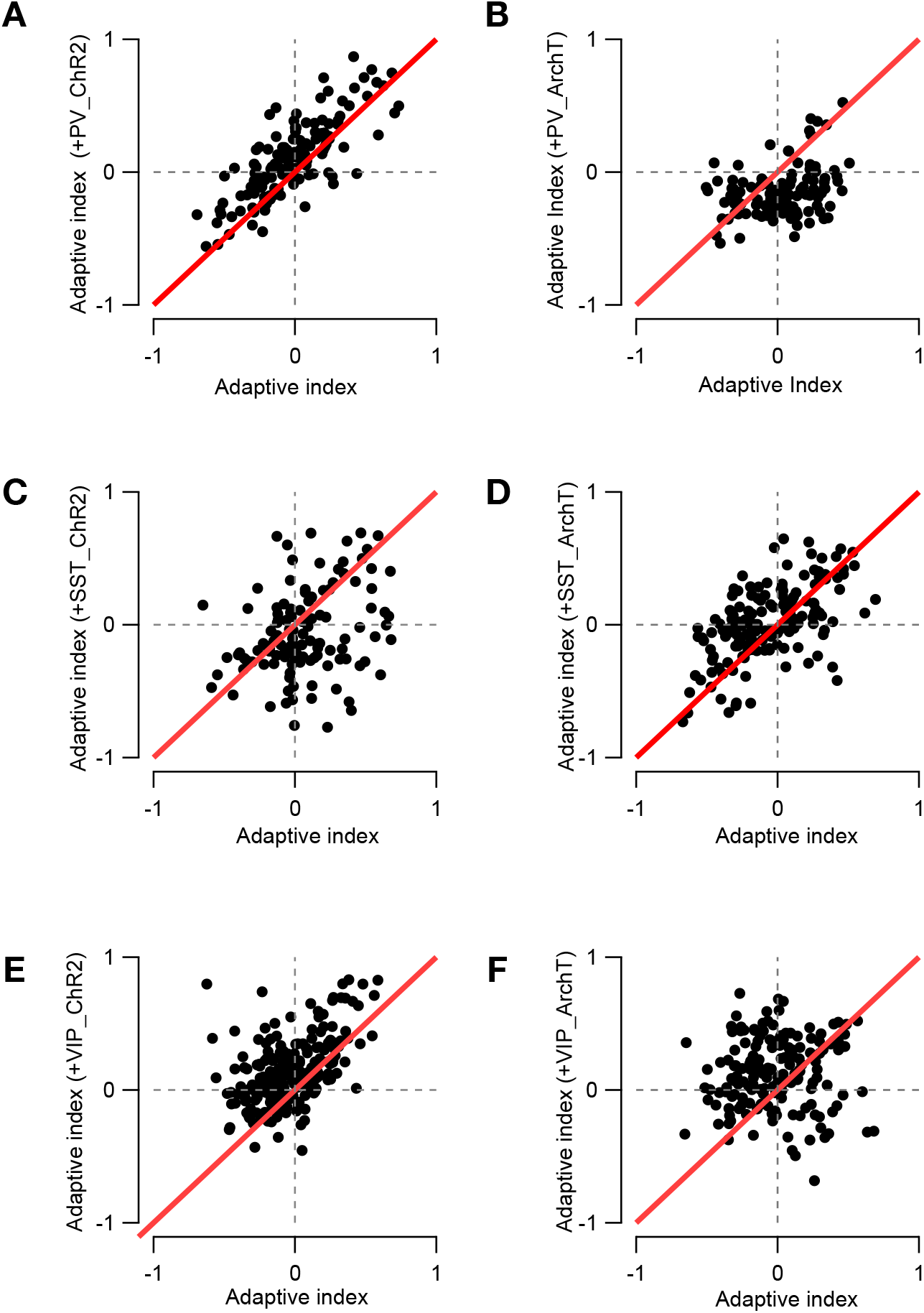
Summary of changes in adaptive indexes of pyramidal cells across conditions of optogenetic manipulations of different interneuron classes. **(A-B)** Scattered plot of Adaptive Index measured from pyramidal cells during optogenetic manipulation of PV interneurons with ChrimsonR (A) and ArchT (B). Abscissa represent AI in control condition and ordinate during optogenetic manipulation of PVs. Negative AI values represent sensitizing-adaptation while positive AI values represent depressing-adaptation). Each black dot represents an individual neuron and the red thick line corresponds to unity. **(C-D)** Similar plot as depicted in A-B but this time during manipulation of SST interneurons. (**E-F**) Similar plot as depicted in A-D but this time during manipulation of VIP interneurons.

**Figure S5.**
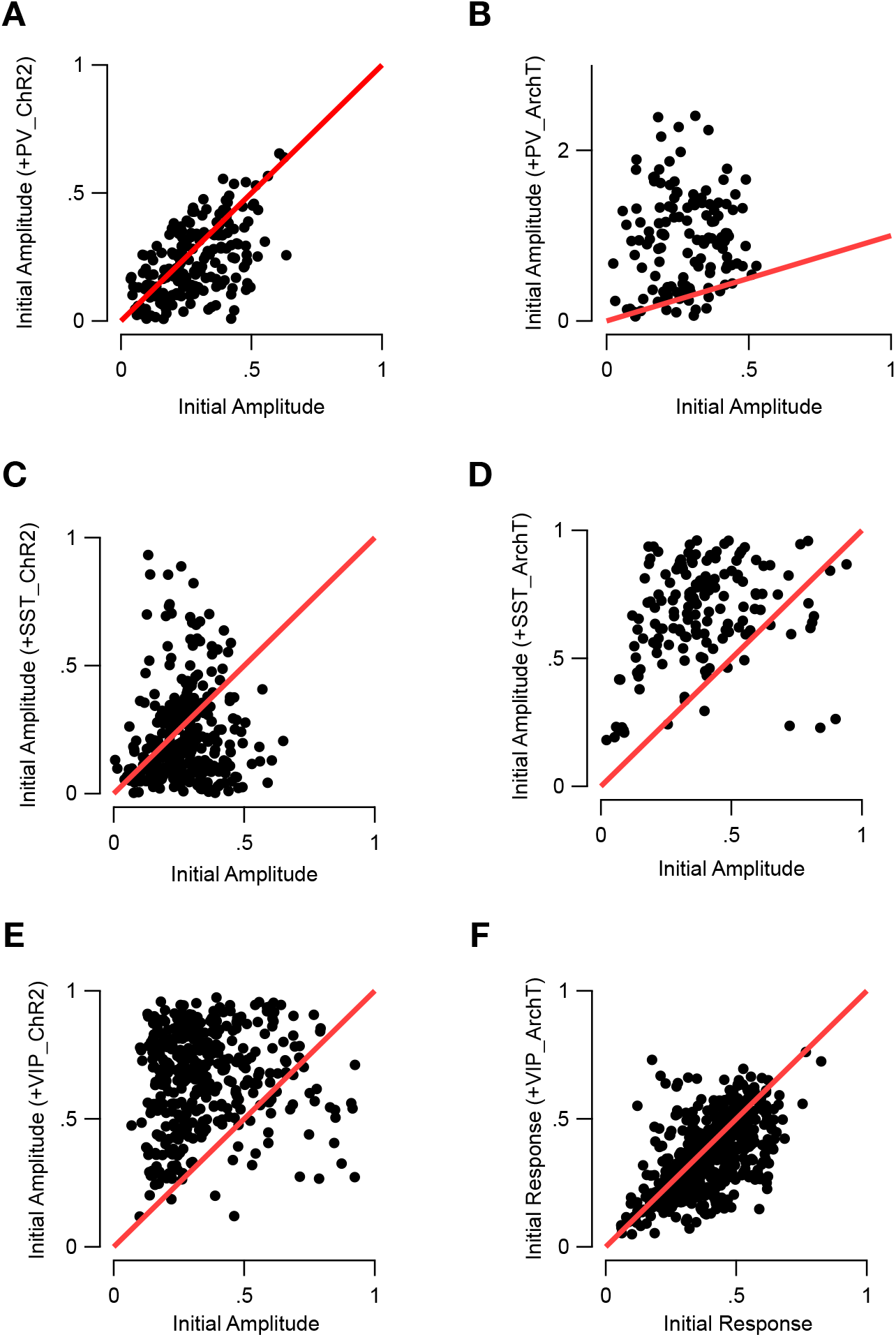
Summary of changes in initial response amplitude of pyramidal cells across conditions of optogenetic manipulations of different interneuron classes. **(A-B)**Scattered plot of initial amplitude of pyramidal cells responses with and without optogenetic manipulation of PV interneurons with either ChrimsonR (A) or ArchT (B). The initial amplitude of responses was measured during the first 2 s of presentation of a high contrast drifting grating. **(C-D)**Similar plot as depicted in A-B but this time during manipulation of SST interneurons. (**E-F**) Similar plot as depicted in A-D but this time during manipulation of VIP interneurons.

